# A Phage Ejectosome Protein Moonlights to Inhibit CBASS Anti-Viral Defense System

**DOI:** 10.1101/2025.07.23.666291

**Authors:** Gong Chen, Ashley Luo, Jacob Bourgeois, Aathmaja Anandhi Rangarajan, Andrew Camilli, Christopher M. Waters, Wai-Leung Ng

## Abstract

A family of prokaryotic antiviral systems collectively known as the <u>c</u>yclic oligonucleotide-<u>ba</u>sed <u>s</u>ignaling <u>s</u>ystem (CBASS) is found in many bacterial species. Phage defense in CBASS is mediated by the action of a specific effector activated by the cognate cyclic oligonucleotide, often resulting in abortive infection. In the prototypical Type II-A CBASS, phage defense is initiated by the production of cyclic GMP-AMP (cGAMP) by enzymes known as cGAS/DncV-like nucleotidyltransferases (CD-NTases) within the bacterial host, and subsequent activation of the CapV phospholipase by cGAMP leads to cell lysis, thereby limiting phage replication within the host population.

Phage proteins that degrade cGAMP, sequester cGAMP, or prevent cGAMP synthesis have been identified in phages to evade CBASS protection. However, some phages are resistant to CBASS protection despite not encoding these proteins, suggesting that additional counter-defense mechanisms exist. To identify new anti-CBASS mechanisms in phages, we exploited bacterial host sensitivity to the antimicrobial compound sulfamethoxazole (SMX) as an indirect read-out for CBASS activity. Bacteria with active CBASS are more sensitive to SMX; therefore, we postulated that any viral proteins produced inside the bacterial host that target CBASS could increase bacterial SMX resistance. Using this screen, we identified that the Gp15 protein from the vibriophage ICP3 increases SMX resistance in *Vibrio cholerae* El Tor biotype that encodes an active CBASS. Gp15 is an essential virion protein involved in DNA ejection and conserved in many phages. Expression of Gp15 of another coliphage increases SMX resistance in *Escherichia coli* expressing an active CBASS, and it allows CBASS-sensitive phage to infect the bacterial host with CBASS, demonstrating an important role of Gp15 in phage defense evasion.

Unlike other known anti-CBASS proteins, Gp15 does not degrade or sequester cGAMP, and it does not reduce the cellular level of cGAMP. Gp15 also does not interfere with CapV binding to cGAMP. Instead, Gp15 directly inhibits cGAMP-activated CapV serine hydrolase activity in a stoichiometric manner. Together, our study reveals a new mechanism for CBASS antagonism, in which a phage DNA ejection protein moonlights to inhibit a CBASS effector, thereby evading anti-phage defense.

## INTRODUCTION

The life cycle and evolution of bacteria are greatly impacted by bacteriophages (phages). Due to strong selective pressure, numerous phage defense systems are present in bacterial hosts [1–10]. Conversely, counter-defense mechanisms are found in phages to evade the protection [11–15]. Recent studies suggest that many eukaryotic anti-viral defense systems have an evolutionary origin from their prokaryotic counterparts [16–23]; therefore, knowledge gained from studying phage-bacteria arms race will inform the impact of phages on bacterial evolution and pathogenesis, and will shed light on the immunity against viral attack in higher organisms.

A family of prokaryotic antiviral systems, collectively known as the <u>c</u>yclic oligonucleotide-<u>ba</u>sed <u>s</u>ignaling <u>s</u>ystem (CBASS), identified in more than 10% of sequenced bacterial species, uses structurally related enzymes known as cGAS/DncV-like nucleotidyltransferases (CD-NTases) to produce a variety of cyclic oligonucleotides, activating various cognate effectors for phage defense[24–27]. In the Type II-A CBASS system in bacteria [24–26, 28, 29], it is postulated that phage infection triggers the production of 3’3’-cyclic GMP-AMP (cGAMP) by the enzyme DncV inside the bacterial host, and subsequent activation of the cognate effector CapV phospholipase by cGAMP leads to cell lysis to limit phage replication within the population, a process known as abortive infection [4, 26, 27]. A related cyclic dinucleotide, 2’3’-cGAMP, made by cGAS (a structural homolog of DncV) stimulates the STING pathway in metazoans for protection against viral infection [30–32]. Thus, cGAMP signaling is considered an evolutionarily conserved pathway for antiviral defense in both prokaryotes and eukaryotes [19, 33, 34].

The current pandemic strain of *V. cholerae,* known as the El Tor biotype, has acquired an operon that encodes a complete Type II-A CBASS, including DncV, CapV, and two other enzymes (Cap2/3) that modulate DncV activity using a ubiquitination-like mechanism [35, 36]. Heterologous expression of this CBASS operon protects *E. coli* from infection by some coliphages [24, 35, 36]. Given its proposed function in antiviral defense, CBASS is believed to confer a fitness advantage to *V. cholerae* El Tor by defending against phage infection. Surprisingly, the three major circulating *V. cholerae* phages (vibriophages) readily evade CBASS to effectively attack El Tor [37–40]. Some coliphages, such as T7, are also notably resistant to CBASS [24, 41]. Three different mechanisms have been identified in phages to evade CBASS protection: Coliphage T4 and some other phages encode a nuclease, Acb1, that degrades cGAMP [42]. In T4 and other *Pseudomonas aeruginosa* phages such as PaMx41, the anti-CBASS protein Vs.4 or Acb2 sequesters cGAMP and other cyclic oligonucleotides to prevent activation of the CBASS effector [36, 43, 44]. From a structure-guided pipeline that predicts potential immune inhibitors, Acb3 was identified to be an inhibitor of the CD-NTases of both Type I and III CBASS [45]. However, based on sequence homology search, the circulating CBASS-resistant vibriophages (ICP1, 2, and 3) do not encode any known anti-CBASS proteins, so it is plausible these phages use a novel mechanism to evade CBASS.

To understand how these vibriophages evade CBASS, we developed an unbiased genetic screen and successfully identified an essential phage protein (Gp15) involved in DNA injection that also moonlights to inhibit CBASS. Our analysis of Gp15 suggests that this protein, unlike other anti-CBASS proteins, targets the cGAMP effector CapV directly. Homologs of Gp15 are found in many phages infecting a variety of bacterial hosts [46–48]; thus, Gp15 may represent a new and conserved mechanism used by phages to evade host protection across different bacterial phyla.

## RESULTS

### A genetic screen to identify novel anti-CBASS phage proteins

The conventional mode of action of the sulfonamide antibiotic sulfamethoxazole (SMX) is to impair folate biosynthesis by inhibiting dihydropteroate synthase. However, folates also function as allosteric inhibitors of cGAMP synthase DncV [49]. Thus, in the presence of SMX, when folate synthesis is inhibited, the activity of DncV increases, resulting in higher cGAMP production and subsequent aberrant activation of CapV, which leads to cell death[50, 51]. Accordingly, we and others have previously demonstrated that *V. cholerae* El Tor with an active CBASS is more sensitive to SMX than mutants lacking a functional CBASS [50, 51].

ICP3, one of the circulating phages predominantly found in cholera patient stools, is related to the prototypical *Podoviridae* phage T7 [37]. Both ICP3 and T7 can infect bacterial hosts carrying CBASS [24, 37, 41], but neither encodes any known anti-CBASS proteins, such as Acb1, Acb2, and Acb3. We therefore hypothesized that these phages use a previously unidentified anti-CBASS mechanism. We first focused on ICP3 because we had previously found that this phage has evolved to escape the defense conferred by two other antiphage systems (DdmABC and AvcID) and infect *V. cholerae* El Tor[52], so it may also have the capability to evade CBASS. We reasoned that any potential anti-CBASS phage protein encoded by ICP3, when produced inside the bacterial cell, could make CBASS-carrying *V. cholerae* El Tor more resistant to SMX, similar to mutants with non-functional CBASS.

We constructed a gene library in which each ICP3 protein-encoding gene (Accession NC_015159.1) was cloned into a plasmid under the transcriptional control of an IPTG-inducible promoter. We introduced this library into *V. cholerae* C6706 (El Tor, CBASS^+^) and monitored the growth of each clone with and without SMX. Some of these ICP3 genes, when induced, inhibited *V. cholerae* growth even without SMX (**Supplementary Table S1**). Out of the remaining ∼50 clones, each expressing a unique non-toxic ICP3 gene, we identified a single clone expressing *gene 15* of ICP3 (YP_004251288.1) that confers higher resistance to SMX (SMX^R^) (**Fig. 1A**). In contrast, expression of *gene 15* does not relieve the growth inhibition due to SMX in a ΔCBASS mutant (**Fig. 1B**), indicating the increase in SMX resistance conferred by *gene 15* expression is CBASS-dependent.

**Fig. 1.**
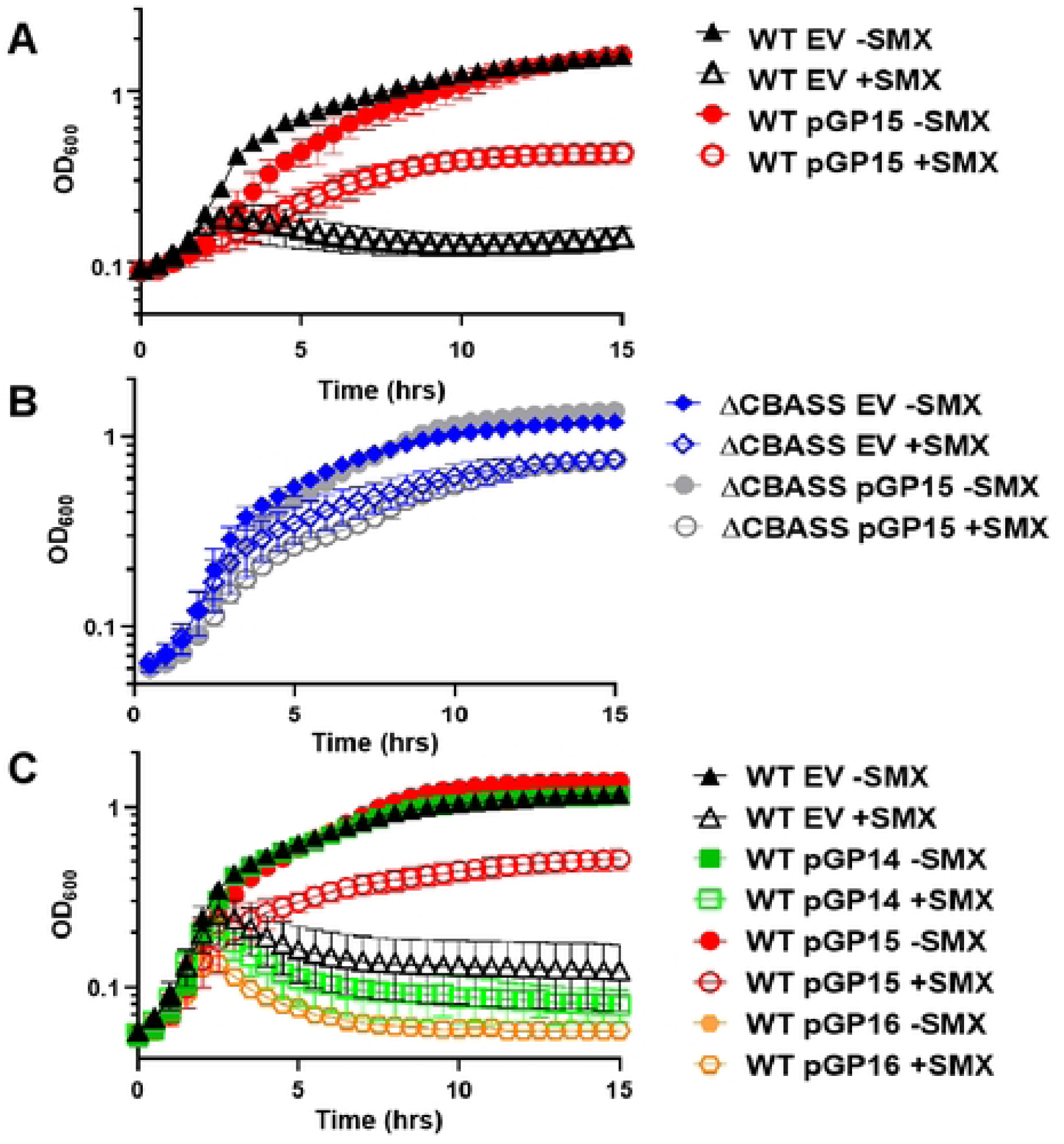
Expression of *gp15* of ICP3 increases sulfamethoxazole (SMX) resistance in *V. cholerae*. (**A**) Growth curves of *V. cholerae* El Tor C6706 (WT) with and without ectopic expression of *gp15* in the presence and absence of 50 µg/mL SMX. (**B**) Growth curves of *V. cholerae* Δ*dncV* Δ*capV* (ΔCBASS) mutants with and without ectopic expression of *gp15* in the presence and absence of 50 µg/mL SMX. (**C**) Growth curves of *V. cholerae* WT with and without ectopic expression of *gp14*, *gp15*, or *gp16* in the presence and absence of 50 µg/mL SMX.

The ICP3 *gene 15* product (Gp15_ICP3_) is homologous to the Gp15 of T7 (NP_042003.1, Gp15_T7,_ E value=9e-89, 65% similarity) [48, 53]. Gp14, Gp15, and Gp16 of T7 form the ejectosome within the virion for phage DNA ejection into host cells [47, 48, 53, 54]. Since only Gp15_ICP3_ was produced in the host cell, the increased SMX^R^ and the potential anti-CBASS activity are not due to the formation of the complete ejectosome. Expression of *gene 14* or *gene 16* of ICP3 alone had no effect in increasing SMX^R^ (**Fig. 1C**). CBASS gene expression was not affected by production of Gp15_ICP3_ (**Supplementary Fig. S1).**

### Conservation and specificity of Gp15 antagonism on CBASS

Phylogenetic analysis shows that homologs of Gp15 are encoded by a broad range of phages that mainly infect Gram-negative bacteria [46, 47] (**Fig. 2A**). To determine if Gp15 from other phages also antagonizes CBASS, we tested two different Gp15 homologs from phages that infect *E. coli*. One is from the canonical phage T7 and the other is from coliphage Mak [55]. These homologs vary in the primary amino acid sequence and length: Gp15_T7_ and Gp15_ICP3_ are similar in length (∼740 amino acids), and Gp15_Mak_ is shorter (692 residues), but Gp15_Mak_ displays a slightly higher similarity to Gp15_ICP3_ than to Gp15_T7_ (**Supplementary Fig. S2A**). Monomeric Gp15_ICP3_ and Gp15_Mak_ predicted by Alphafold3 [56] are both structurally similar to the published structure of Gp15_T7_ determined by Cryo-EM [47, 54] (**Fig. 2B**). We first verified that an *E. coli* strain without a native CBASS expressing the Type II-A CBASS operon from *E. coli* TW11681 [24] on a plasmid is more SMX sensitive than the empty vector control (**Fig. 2C**). Production of Gp15_Mak,_ but not Gp15_T7,_ relieved the SMX toxicity caused by CBASS_Ec_ in this strain (**Fig. 2C**). For unknown reason, production of Gp15_ICP3_ in the presence of SMX, even without CBASS, was toxic to the *E. coli* host and therefore was not followed up.

**Fig. 2.**
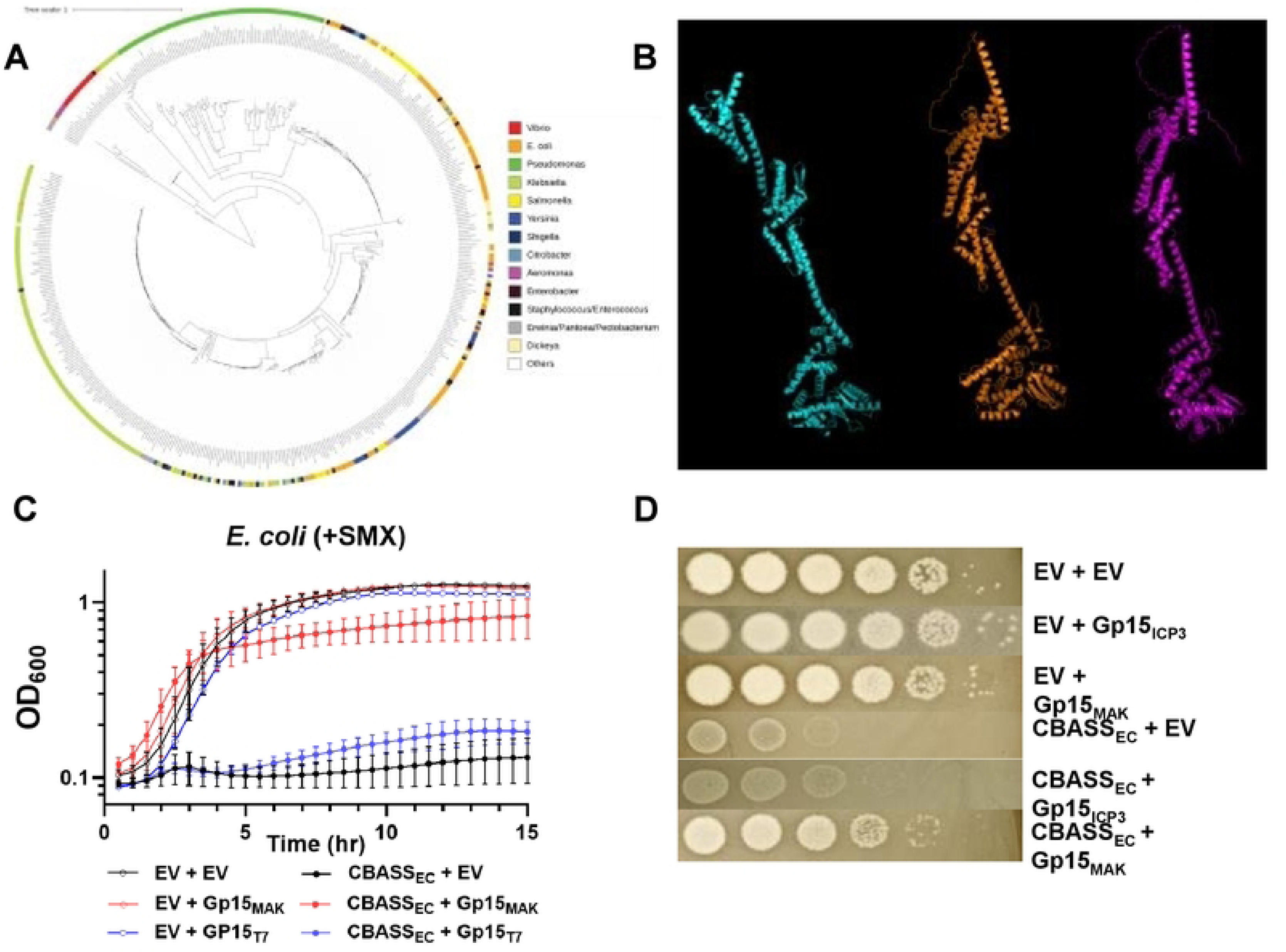
Conservation and specificity of Gp15 antagonism of CBASS. (A) Homologs of Gp15_ICP3_ (YP_004251288.1) were identified by searching against the NCBI protein database using BLASTP. Over 400 hits were identified, and their sequences were retrieved (cutoff E value <1E-20 and coverage >70%). The phylogenetic tree of these Gp15 homologs is built with COBLAT^26^. (B) Structural homology between Gp15_T7_ (left, cyan, PDB 7k5c_A,), Gp15_Mak_ (middle, orange, predicted by Alphafold 3), and Gp15_ICP3_ (right, magenta, predicted by Alphafold 3). Structural alignment was performed by PyMOL. (**C**) Growth of *E. coli* strains in the presence of 50 µg/mL SMX carrying a plasmid with and without the CBASS operon from *E. coli* TW11681, and in the absence and presence of Gp15 from T7 or coliphage Mak. (**D**) T2 infection efficiency on different *E. coli* strains with and without CBASS and GP15_ICP3_ or Gp15_MAK_. The experiment was repeated more than three times, and representative graphs are shown.

Since SMX sensitivity is only an indirect readout of CBASS, we investigated whether Gp15 plays a direct role in antagonizing CBASS during phage infection. However, since Gp15_T7_ is essential for T7 [48] and Gp15_ICP3_ is likely also essential, we did not attempt to construct an ICP3 phage mutant lacking Gp15. Moreover, no vibriophage has been identified as CBASS-sensitive, so we could not directly test the anti-CBASS function of Gp15_ICP3_ in *V. cholerae*. Instead, we tested whether Gp15 from different phages inhibits CBASS_Ec,_ allowing the infection of a CBASS-sensitive phage that does not encode Gp15. As shown before [24, 35] and here, the production of CBASS_Ec_ in *E. coli* reduced phage T2 infection efficiency by ∼3-4 orders of magnitude (**Fig. 2D**). In contrast, T2 plaquing efficiency was partially and nearly restored when Gp15_ICP3_ or Gp15_mak_ was co-produced in cells with CBASS_Ec_, respectively (**Fig. 2D**). Consistent with the lack of effect of Gp15_T7_ in the SMX assay, production of Gp15_T7_ did not restore T2 infection in the presence of CBASS_EC_ (**Supplementary Fig. S2B**). Together, our results suggest that Gp15 from multiple phages exhibits anti-CBASS activity and can be used to evade CBASS during phage infection. While CBASS antagonism by Gp15 is conserved in some phages, specificity among different phage-host pairs is observed.

### Gp15 does not affect cGAMP levellevels in *V. cholerae*

Known anti-CBASS proteins, including Acb1, Acb2, and Acb3, use different mechanisms to reduce the level of cGAMP available to activate the cognate effector to evade the phage defense [36, 42–45]. Therefore, we first tested if Gp15_ICP3_ reduces cGAMP levels inside *V. cholerae* cells. To allow robust detection of cGAMP and avoid growth inhibition due to cGAMP overproduction [28], we overexpressed *dncV* in *V. cholerae* without CapV and used LC-MS/MS [51] to determine the concentration of cGAMP inside the cell. We detected a significant increase in cGAMP production when *dncV* was overexpressed, but the cGAMP level inside the cell was not significantly reduced with co-expression of Gp15_ICP3_, suggesting Gp15 does not inhibit the activity of DncV, and it does not actively degrade cGAMP (**Fig. 3A**).

**Fig. 3.**
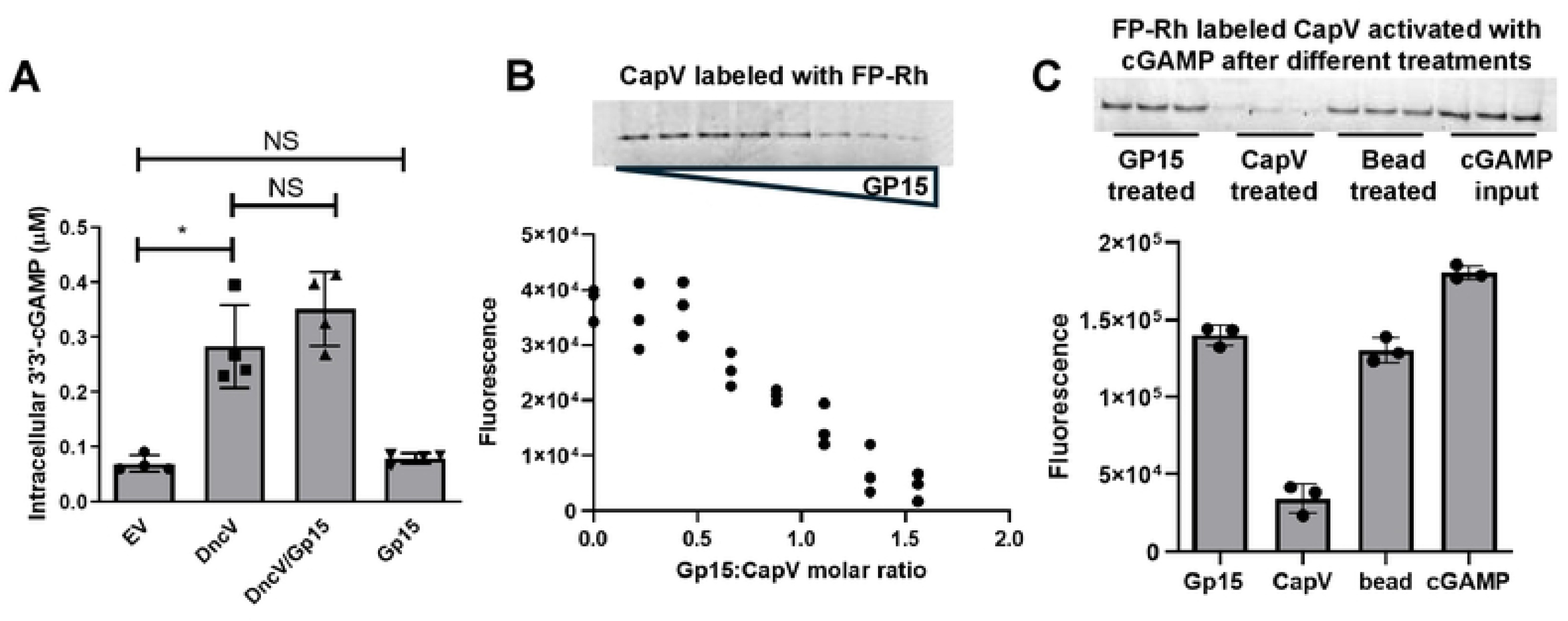
GP15 inhibits CapV hydrolase activity. **(A)** Intracellular concentration of cGAMP in *V. cholerae* Δ*dncV* Δ*capV* mutants with and without ectopically expressing *dncV* and with and without co-expressing *gp15* of ICP3. cGAMP levels in the samples were determined by LC-MS/MS using a cGAMP standard curve and normalized to the viable cell counts. A pairwise comparison was performed with a *t*-test. (* *p*=0.0286). (**B**) Top: CapV activation level determined by FP-Rh labeling in the presence of increasing concentrations of Gp15. Bottom: The fluorescent signals were quantified and plotted against the ratio between CapV and Gp15. In each reaction, both CapV and cGAMP were used at 0.7 µM, and Gp15 was used from 0 to 1.1 µM. (**C**) Top: CapV activation level determined by FP-Rh labeling in the presence of cGAMP after different treatments. The input cGAMP was pretreated with Gp15 or CapV, and the proteins were removed by Ni-NTA beads. Bottom: The fluorescent signals were quantified. In all cases, each experiment was repeated three times, and representative results are shown with multiple technical replicates (n ≥ 3).

### Gp15 inhibits cGAMP-activated CapV

We then tested if Gp15 directly inhibits CapV hydrolase activity. CapV activity was tested using a serine hydrolase assay we previously developed [28]. In this assay, purified cGAMP activates CapV, and the activated CapV can be detected by covalent labeling with a rhodamine-labeled fluorophosphonate probe (**FP-Rh**) [28]. In the presence of increasing amounts (0 to 1.5 µM) of purified Gp15_ICP3_, CapV_Vc_ (0.7 µM) activation by cGAMP was inhibited when the molar ratio of Gp15_ICP3_ and CapV_Vc_ is ≥ 1:1 (**Fig. 3B**), indicating Gp15_ICP3_ directly interferes with CapV_Vc,_ likely with a non-enzymatic mechanism.

### Gp15 does not bind or degrade cGAMP *in vitro*

As shown above, Gp15 does not reduce cGAMP levels inside bacterial cells, but it inhibits CapV. To further test if Gp15 sequesters cGAMP with a mechanism similar to Acb2, cGAMP was mixed with a 10-fold excess of purified His_6_-tagged Gp15_ICP3_ or CapV_Vc_ immobilized onto Ni-NTA beads; after incubation, the beads containing the proteins and any bound cGAMP were removed, and the remaining cGAMP in solution was tested using a CapV activation assay as above. As expected, CapV_Vc_ effectively captured most of the cGAMP from the solution, and there was not much cGAMP remaining in the solution to activate CapV_Vc_ in the labeling assay, as shown by the weak FP-Rh signal (**Fig. 3C**). In contrast, Gp15_ICP3_ did not significantly remove or degrade any cGAMP under the same conditions, as shown by the strong FP-Rh labeling signal, which was similar to the signal from the bead-only negative control, and the signals were only slightly lower than the cGAMP input (**Fig. 3C**). Together, our results indicate that Gp15 does not directly target cGAMP to inhibit CapV.

### Gp15 does not prevent CapV from binding to cGAMP

While Gp15_ICP3_ does not bind cGAMP effectively, we tested if the phage protein prevents CapV from binding to cGAMP as a potential CBASS inhibition mechanism. cGAMP was mixed with CapV in the presence of either Gp15_ICP3_ or BSA, or just with just Gp15_ICP3_ or BSA alone. After incubation, each mixture was filtered through a membrane with a MW cutoff of 10 KDa to remove the proteins and any bound cGAMP; the unbound cGAMP in the filtrate was then measured by the CapV activation assay or by LC-MS/MS as described [51]. We use a membrane filter instead of Ni^2+^-NTA beads, as described above, because both CapV and Gp15 are His_6_-tagged; thus, using beads may prevent the interaction of the two proteins. As expected, from these assays, we did not detect any significant binding of cGAMP with Gp15_ICP3_ alone nor with BSA, and the concentration of the unbound cGAMP in the filtrate was similar to the no protein control (**Fig. 4A-B**). In contrast, regardless of whether Gp15 or BSA was present, CapV_Vc_ effectively captured cGAMP in solution, reducing the concentration of cGAMP in the filtrate (**Fig. 4A-B**). The concentrations of cGAMP in the filtrate solutions determined by LC-MS/MS matched closely to the Rh-FP labeling results (**Fig. 4A-B**). These results suggest that CapV binding to cGAMP is not inhibited by Gp15.

**Fig. 4.**
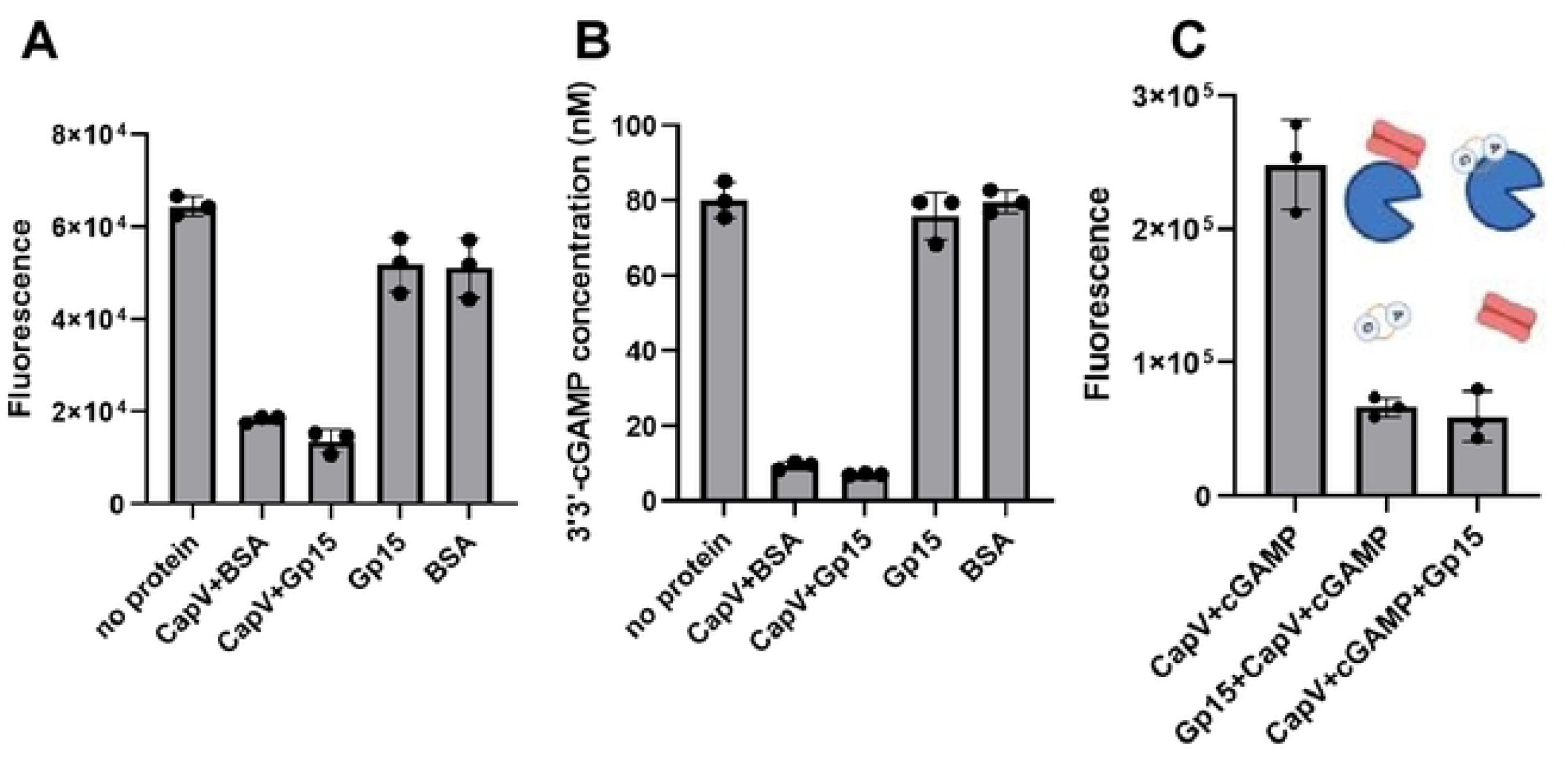
GP15 inhibits CapV not by targeting cGAMP. **(A)** CapV activation level determined by FP-Rh labeling in the presence of cGAMP after different treatments. The input cGAMP was either untreated (no protein) or pretreated with Gp15 alone, BSA alone, CapV and BSA together, or CapV and Gp15 together, and the proteins were removed by filtering through a 10 KDa MWCO filter. The fluorescent signals of the gel were then quantified. (**B**) The cGAMP levels in the samples shown in (A) were quantified by LC-MS/MS. (**C**) CapV activation level determined by FP-Rh labeling. CapV was activated by cGAMP without Gp15 (left), or CapV was activated by cGAMP after addition of Gp15 (middle), or CapV was activated by cGAMP before the addition of Gp15 (right). Each experiment was repeated three times, and representative results are shown with multiple technical replicates (n ≥ 3).

We then tested if Gp15_ICP3_ can inhibit CapV even after it is activated by cGAMP. As shown in the *in vitro* assays above, the interaction between cGAMP and CapV_Vc_ is relatively stable in solution (**Fig. 3C and 4A-B**). Therefore, we first activated CapV_Vc_ with cGAMP, then added purified Gp15_ICP3_ to the reaction, and stopped it with Rh-FP labeling. We observed inhibition on CapV_Vc_ by Gp15_ICP3_ regardless of whether cGAMP was added before or after Gp15_ICP3_ addition (**Fig. 4C**), suggesting Gp15 acts on CapV even if cGAMP already activates it in the cell.

### Gp15 directly interacts with CapV inside the bacterial cell

Our *in vitro* assays shown above indicate that Gp15 directly inhibits the serine hydrolase activity of CapV. Therefore, we used a bacterial two-hybrid system (**BACTH**) [57] to test whether these two proteins interact directly inside bacterial cells. We constructed fusions of Gp15 and CapV to the complementary fragments of the *Bordetella pertussis* adenylate cyclase, known as T25 and T18, in *E. coli*. If these two proteins interact within the cell, functional adenylate cyclase will form, leading to cAMP synthesis and activation of the reporter *lacZ* gene, which can be detected as blue colonies on LB plates with an X-gal indicator. As expected, blue colonies were obtained when Gp15-T25 and Gp15-T18 were co-produced since Gp15 forms oligomers [47, 48, 54]. When CapV-T25 and Gp15-T18 were co-expressed, we detected blue colonies, but not in empty vector negative controls (**Fig. 5A**), indicating that Gp15 and CapV interact directly. These interactions were further quantified using β-galactosidase assays, and the results confirmed that CapV and Gp15 interact within bacterial cells (**Fig. 5B**).

**Fig. 5.**
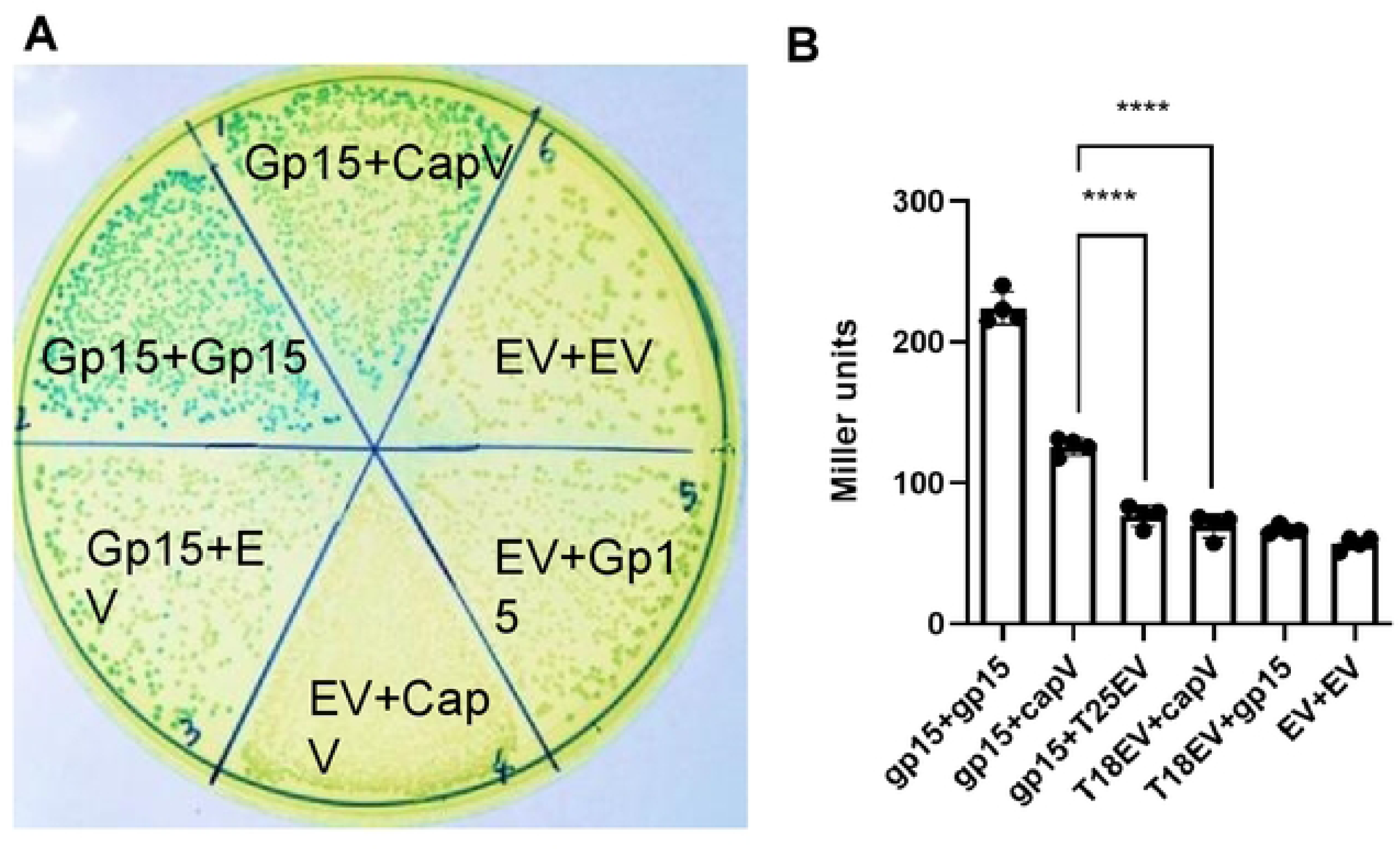
Gp15 and CapV directly interact within bacterial cells. (**A**). The BACTH system was used to study the interaction between Gp15 and CapV. *E. coli* strains co-expressing different Gp15 and CapV T18/T25 fusion proteins were plated on a medium containing X-Gal. EV denotes an empty vector. (**B**) Quantitation of the level of β-galactosidase in different strains co-expressing different CapV or Gp15 fusions. Results were analyzed by ANOVA (\*\*\*\**p* < 0.0001); the assays were repeated twice, and representative results with multiple technical replicates are shown (mean ± SD).

### Gp15 level is higher than CapV inside *V. cholerae* cells during phage infection

Our analyses above suggest that Gp15 inhibits CapV with a non-enzymatic mechanism, and the phage protein must be present at least in an equimolar ratio with the effector to be effective. To measure the levels of these proteins in cells, we infected *V. cholerae* El Tor with ICP3 at an MOI of ∼5 to ensure that almost every cell was infected, and cell lysates were prepared at different time points during phage infection (0 to 25 minutes). Newly synthesized Gp15 was detected in infected *V. cholerae* cells via Western blot as early as 5 minutes after phage infection (**Fig. 6A, Supplementary Fig. S3A**), and the level of Gp15 increased steadily from 5 to 15 minutes, followed by cell lysis. Using various amounts of purified Gp15_ICP3_ mixed with *V. cholerae* cell lysates as standards, we used quantitative Western blotting to determine that Gp15_ICP3_ is present at ∼2500-10,000 copies per *V. cholerae* cell during phage infection (**Fig. 6B, Supplementary Fig. S3B**). In contrast, the level of CapV_Vc_ in wild type was below the limit of detection by Western blot under the same conditions. Based on a standard curve generated using purified CapV_Vc_ mixed with cell lysates prepared from the Δ*capV* mutant, it was estimated to be below 200 copies per cell (**Fig. 6C, Supplementary Fig. S3B**). In contrast, when we overexpressed CapV inside *V. cholerae*, we could readily detect the protein using the same conditions (**Fig. 6C**). Although undetectable by our Western blot assays, CapV must be present and active inside *V. cholerae* El Tor because the wild-type bacterium is more sensitive to SMX than its Δ*capV* mutant [50, 51], and overexpression of *dncV* alone induces CapV-dependent cell lysis [28]. The burst size of ICP3 is ∼50 phages per cell [52], and each T7 viral particle contains eight copies of Gp15 [58]. Based on our quantitation results and assuming ICP3 also contains eight copies of Gp15 in each phage particle, we conclude that even though a significant portion of Gp15 is packaged into virions, the amount of unincorporated Gp15 produced during phage infection is still in excess of CapV inside infected *V. cholerae* cells, sufficient to inhibit CapV to evade CBASS protection. Together, our results strongly suggest Gp15 moonlights as a new anti-CBASS protein in addition to its role in DNA ejection.

**Fig 6.**
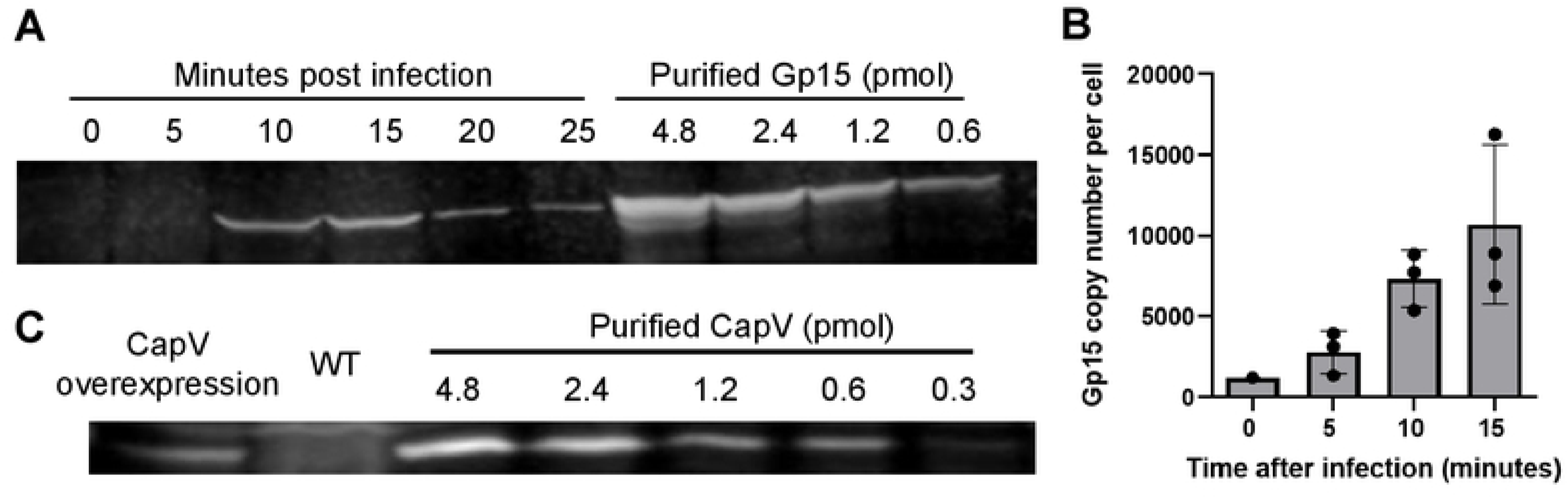
Gp15 cellular level is in excess of CapV during phage infection. **(A)** Western blot using anti-Gp15 antibodies with cell lysates prepared at different time points after ICP3 infection of *V. cholerae* WT. The Gp15 standard curve was prepared by mixing different amounts of purified His_6_-Gp15 with *V. cholerae* WT cell lysate without infection. (**B**) Quantitation of Gp15 amount per cell during ICP3 phage infection. (**C**) Western blot using anti-CapV antibodies with cell lysates from *V. cholerae* wild type and a strain ectopically overexpressing CapV. The CapV standard curve was prepared by mixing different amounts of purified His_6_-CapV with cell lysates prepared from a strain lacking CapV.

## Discussion

Multiple phage defense systems, including CBASS, DdmABC, and AvcID, are encoded by the two genomic islands (VSP-1 and VSP-2) in the current 7^th^ pandemic strain of *V. cholerae* El Tor biotype [52, 59–62]. The acquisition of these two genomic islands could potentially enhance the fitness of El Tor biotypes by protecting against infection by circulating bacteriophages. Yet, the three different vibriophages (ICP1, ICP2, and ICP3) commonly found in cholera patients readily infect *V. cholerae* El Tor biotype, and how these phages evade the protection conferred by these antiphage defense systems on VSP1 and VSP-2 remains unclear.

We have previously identified that phage DNA polymerase polymorphism in different ICP3 phage isolates contributes to the sensitivity of DdmABC and AvcID [52]. Here we show that Gp15 from ICP3 can directly target the CBASS effector CapV phospholipase using a mechanism distinct from the other anti-CBASS proteins that act on cGAMP level or synthesis [36, 42–45]. The ability to directly antagonize CapV by Gp15 appears to be conserved in other phages, such as coliphage Mak, as revealed in this study. To our knowledge, this is the first example of an anti-CBASS protein that does not reduce the level of cGAMP available to activate the cognate CBASS effector. The advantage of targeting the effector is over targeting the cGAMP messenger directly is unclear, but protein-protein interaction likely offers higher antagonism specificity and stringency. In the case of cGAMP-targeting anti-CBASS proteins, specificity appears to be more relaxed. For example, Acb1 from T4 has a broad cyclic nucleotide hydrolysis activity and degrades cGAMP and some other related cyclic oligonucleotides [42], and Acb2 proteins from T4 and *Pseudomonas* phage PaMx33 display high affinity towards multiple cyclic trinucleotides and dinucleotides [44]. Acb3 is thought to target CD-NTases through direct protein-protein interaction; however, the phage protein exhibits relaxed specificity, capable of inhibiting both Type I and Type III CBASS, as well as the eukaryotic cGAS enzyme[45] [45]. In our case, the antagonism of Gp15 toward CapV seems to be phage-host pair specific, as Gp15 from ICP3 and Mak, but not T7, antagonize Type II-A CBASS. This specificity could be due to the co-evolution of the interacting partners, resulting in a unique interaction surface between the phage protein and the host enzyme. The 3D structure of T7 Gp15 has been solved [47, 54], and the structures of the apo form and the cUA-bound form of CapE (a homolog of CapV) have been recently experimentally determined [63]. We have attempted to use Alphafold 3 [56] to predict the complex formed by Gp15 and CapV; however, the resulting Alphafold 3 model has a low confidence score, providing insufficient information to determine the potential interacting surface. The inability to generate a reliable structural model for the complex using the current Alphafold algorithm could be due to the significant conformational or other physical changes such as filamentation in the cGAMP-activated CapV [63]. Additional genetic and structural analyses are necessary to identify the interacting surfaces between CapV and Gp15.

The other unique feature of Gp15 is its moonlighting properties. Moonlighting proteins carry multiple biochemical or biophysical functions within a single peptide chain that do not arise due to gene fusion, and these functions are physiologically distinct [64, 65]. In viruses that infect eukaryotes, viral surface glycoproteins that carry multiple unique functions are not uncommon, and some of these proteins are used for immune evasion [66]. Bacteriophages also use moonlight proteins to carry out multiple processes during phage infection, often to facilitate their propagation [67, 68]. The common use of moonlight proteins for viruses likely allows the maintenance of the relatively small and streamlined genomes that can be packaged effectively within the capsid without errors. Gp15 is part of the ejectosome of the phage T7, and it is expected that Gp15 performs the same role in ICP3 due to the high sequence homology between the two proteins. We postulate that Gp15 is utilized in multiple steps during phage replication. The Gp15 carried inside the virion is involved in DNA ejection after host cell attachment, while the Gp15 produced in the cytoplasm during phage replication is used for both virion assembly and targeting CapV. Therefore, the timing of Gp15 synthesis, assembly into the virion, and mounting the counter-defense must be precisely coordinated. ICP3 has evolved to solve this problem by producing a large amount of Gp15 at a relatively early stage during phage replication. The amount of Gp15 in infected *V. cholerae* cells exceeds that of CapV, ensuring a stoichiometric interaction between the two proteins without affecting virion assembly.

While the current study illustrates a new anti-CBASS mechanism employed by some phages, it remains unclear how other phages that do not encode any cGAMP or CapV targeting proteins evade CBASS protection. For example, ICP1 and ICP2 can infect El Tor effectively, but they do not encode homologs of any anti-CBASS proteins. As folate depletion could potentially act as a cue for CBASS activation during phage infection [49–51], some CBASS-resistant phages may encode functions to maintain the level of folate within the infected bacterial hosts. Studies of escape phages resistant to CBASS and protein-protein interaction analysis revealed that phage capsid proteins could serve as triggers to activate different CBASS systems [43, 69, 70]. Therefore, ICP1 and ICP2 may have evolved to encode capsid proteins that enable these phages to evade the surveillance mechanism of CBASS, although capsid protein activation of *V. cholerae* CBASS has not been demonstrated. These phages may also encode novel proteins to circumvent CBASS to evade immunity.

The phage-bacteria arms race often drives rapid evolution in the genomes of both the bacterial hosts and the infecting phages, resulting in the loss and gain of genetic content over time. Therefore, it remains an enigma that the circulating El Tor strains, especially those isolated from cholera patients, retain the possession of VSP-1 and VSP-2, despite these islands being ineffective in protecting against phage infection. For VSP-2, the DdmABC system may confer fitness for eliminating plasmids or other foreign DNA [60, 62]. Previous studies also indicate that VSP-1 and VSP-2 in the El Tor biotype could provide certain unknown metabolic functions that could help to outcompete the extinct classical biotype [71]. VSP-2 also encodes functions important for adaptation to zinc starvation [72]. Thus, it is plausible that CBASS and other systems encoded in these islands are involved in other processes that provide fitness advantages to the El Tor biotypes, independent of their role in phage defense. Finally, it is also possible that circulating phages that are sensitive to these anti-phage systems have yet to be identified. For example, we have previously identified a rare circulating ICP3 isolate in a cholera patient’s stool sample that is sensitive to both DdmABC and AvcID [52], so it is likely that some vibriophages that have yet to be discovered are sensitive to CBASS. Further studies on the physiological functions of CBASS and other genes in the VSP islands, as well as phages that overcome the defense conferred by these islands, will shed light on the evolution of this deadly pathogen.

## Materials and Methods

### Bacterial strains, phages, and plasmids

*V*. *cholerae* strains used in this study were derived from C6706str2, a streptomycin-resistant isolate of C6706 (O1 El Tor) [73] or from E7946 [74]. A complete list of bacterial strains and plasmids used in this study is provided in Supplementary Table S2. Primers used in this study is provided in Supplementary Table S3. *V*. *cholerae* and *E*. *coli* cultures were grown with aeration in Lysogeny Broth (LB). Unless specified, media was supplemented with streptomycin (Sm, 100 μg/ml), Carbenicillin (Carb, 50 μg/ml), kanamycin (Kan, 100 μg/ml), chloramphenicol (Cm, 2.5 μg/ml) and polymyxin B (Pb, 50 U/ml) when appropriate. Phage propagation, phage infection, and phage titer determination were performed as previously described [52]. The ORF of T7 Gp15 (NP_042003.1) was PCR amplified from the genomic DNA prepared from phage T7, and Genewiz synthesized the ORF of Mak Gp15 (CAH6634586.1). These Gp15 coding sequences were then individually cloned into pMMBNeo, a kanamycin-resistant derivative of broad-host range plasmid pMMB67eh [75].

### Construction of the ICP3 open reading frame (ORF) library

ICP3 ORFs (NCBI Accession NC_015159.1) were individually PCR amplified from genomic DNA using primers that added 5’ PstI and 3’ SalI restriction sites. To standardize translation efficiency between ORFs, primers included a strong RBS, 5’-TTTAGGATACATTTTT-3’, directly upstream of the start codon of each ORF [76]. The PCR products were digested with PstI and SalI and cloned into the expression vector pMMBneo. Ligation products were first transformed into *E. coli* TG1. Plasmids containing the ICP3 ORFs were identified from TG1 transformants and transformed into *E. coli* MFDpir [77] and subsequently conjugated into *E. coli* MG1655. Each plasmid of the final ICP3 ORF library was verified by whole plasmid sequencing.

### Genetic screen to identify putative anti-CBASS gene in ICP3

Each plasmid of the ICP3 ORF library constructed in *E. coli* MG1655 was introduced into WT *V. cholerae* C6706 through tri-parental mating using a modified version (Cm resistant) of helper plasmid pRK2013. Kanamycin-resistant transconjugants from each mating were picked and resuspended in 50 µl LB. Ten µl of the cell suspension was added to 90 µl LB with 50 µg/ml Kan, 1 µM IPTG, and with or without 50 µg/ml Sulfamethoxazole (SMX) dissolved in DMSO in each well of a transparent 384-well plate. The cultures were incubated at 30°C under constant shaking, and OD_600_ was monitored every 30 minutes for 20 hours by a Synergy™ HTX Microplate Reader. The growth curves were then compared to that of *V. cholerae* harboring a pMMBneo empty vector. Putative SMX-resistant clones were further tested using the assay described below.

### Sulfamethoxazole (SMX) resistance assay

To assess SMX resistance of different *V. cholerae* and *E. coli* strains, three isolated colonies from each strain were resuspended in 50 µL of LB. Ten µL of the suspension was added to each well of a transparent 96-well microplate that contains 150 µL of LB with or without 50 µg/mL SMX and 1 mM IPTG for *V. cholerae* or 0.1 mM IPTG for *E. coli*. For clones expressing Gp15 from pMMBNeo, 100 µg/mL kanamycin was included. For clones expressing the Type-II *E. coli* CBASS system from pLOCO2 [35] (a gift from Dr. Aaron Whiteley), 50 µg/mL of carbenicillin was included. For *V. cholerae* cultures, each well was covered with 40 µl mineral oil, and a lid was not used. For *E. coli* culture, mineral oil was not used due to aggregate formation; instead, a lid treated with 0.1% Triton X-100 in 100% ethanol was used to reduce condensation and prevent evaporation. The cultures were incubated at 30°C (*V. cholerae*) or 37°C for 15 hours (*E. coli*) under constant shaking, and OD_600_ was monitored every 30 minutes by a microplate reader.

### T2 infection assay in *E. coli*

T2 infection assays were performed using *E. coli* MG1655 containing different plasmids with and without CBASS and different Gp15. Strains were grown in LB containing 100 µg/mL kanamycin and 50 µg/mL carbenicillin overnight and then diluted to an OD_600 of_ ∼0.05 in the same medium with 0.1 mM IPTG. They were subsequently grown to an OD_600 of_ ∼0.5. One hundred µL of each culture was added to 6 mL of 0.4% LB soft agar with 100 µg/mL kanamycin, 50 µg/mL carbenicillin, and 0.1 mM IPTG, and the mixture was immediately poured onto an empty petri plate. After drying for 15 minutes, 10 µL of each T2 serial dilution was spotted onto each plate and left for 20 minutes until visibly dry. Plaques formed on the lawn of bacteria were imaged after overnight incubation at 37°C.

### Bioinformatic analysis of Gp15 in different phages

Homologs of Gp15 of ICP3 (YP_004251288.1) were identified by searching against the NCBI protein database using BLASTP (cutoff E value <1E-20 and coverage >70%). The sequences of more than 400 hits were retrieved and the phylogenetic tree of these Gp15 homologs was built with COBLAT [78]. The 3D structure of Gp15 of ICP3 or Mak was predicted using Alphafold 3 [56]. The predicted structure was aligned with the Gp15 of T7 using PyMOL 3.0.

### Recombinant protein purification (CapV and Gp15)

Recombinant CapV_Vc_-His_6_ was purified as described previously without any modifications [28]. For recombinant Gp15_ICP3_ purification, a protocol for purifying recombinant Gp15_T7_ from *E. coli* was used with modifications [48]. *E. coli* BL21(DE3) carrying pET28b-His_6_-Gp15_ICP3_ was grown overnight in LB with 50 µg/mL Kanamycin; the overnight culture was then diluted 100-fold in 1 L of the same medium and grown at 37°C until OD_600_ reached 0.5. Protein expression was induced with 1 mM IPTG at 16°C overnight. Cells were harvested by centrifugation and then resuspended in 20 ml lysis buffer (50 mM HEPES-KOH pH 8.0, 300 mM KCl, 20% (v/v) glycerol, 1 mM betaine, 10 mM imidazole) and lysed with a Microfluidizer^®^ M-110S. The lysate was centrifuged at 4°C, 10,000g for 1 h to pellet cell debris. The supernatant was filtered with a 0.45 µM syringe filter (Millipore) and then loaded onto a 1 mL HisTrap^TM^ HP Prepacked column (GE Healthcare). The column was washed sequentially with 10 mL lysis buffer with 20 mM and 50 mM imidazole. Proteins were eluted with 10 mL elution buffer (lysis buffer supplemented with 50 mM potassium glutamate, 50 mM arginine, 10 mM MgCl_2_, 10 mM CaCl_2_, 300 mM imidazole) and collected in 500 µl fractions. After analysis with SDS-PAGE gel, fractions with the least contamination and most Gp15 were pooled and dialyzed using a 10 K MWCO Slide-A-Lyzer dialysis cassette (Thermofisher) in dialysis buffer (50 mM HEPES-KOH pH 8.0, 300 mM KCl, 1 mM betaine). Dialysis was performed at 4°C in 1 L buffer, which was changed twice at 2-hour intervals, followed by a final overnight dialysis. The resulting protein was quantified with Bio-Rad Protein Assay reagent and stored in 50 µL aliquots at −80°C before use.

### CapV activation assay

Purified CapV_Vc_-His_6_ was first diluted in FP-Rh buffer (50 mM sodium phosphate (pH 7.4), 300 mM NaCl, 10% (v/v) glycerol) to a final concentration of 9.5 µM. The CapV activation assay contains 1 µl of diluted CapV_Vc_-His_6_, 0 to 7 µl His_6_-Gp15_ICP3_ (2.1 µM, in 1 µl increments), 7 to 0 µl Gp15 dialysis buffer, 1 µl 3’3’-cGAMP (10 µM in water, InvivoGen), and 4 µl FP-Rh buffer. The reaction was incubated for 20 min at room temperature. Subsequently, 1 µl fluorophosphonate-rhodamine probe (FP-Rh, 5 µM in DMSO, a generous gift from Dr. Aimee Shen, Tufts University) was added to the mixture and incubated for 30 min in the dark. The reaction was stopped by adding 4.7 µl of 4x SDS loading dye and 15 µl of the final mixture was loaded onto a 10% SDS-PAGE gel. After electrophoresis, gels were scanned using a Fujifilm Starion FLA-9000 image scanner with 532 nm excitation and BPG1 (570DF20) emission filter, and the fluorescence intensity was quantified using ImageJ software.

### cGAMP binding assay

To determine cGAMP binding by proteins, His_6_-tagged CapV or Gp15 was first captured on Ni-NTA magnetic agarose beads (Pierce). For each sample, 10 µl of the beads was first washed twice with 50 µl of Gp15 dialysis buffer, and then 100 pmol of His_6_-tagged CapV or Gp15 (in Gp15 dialysis buffer with 0.05% Tween-20) was added to the washed beads and incubated at room temperature for 20 minutes with constant mixing. Washed beads were incubated with 50 µl Gp15 dialysis buffer as controls. Any unbound proteins were then removed, and the beads were washed again twice with 50 µl of Gp15 dialysis buffer. Then, 10 µl of cGAMP (1 µM in water) was incubated with the beads at room temperature for 10 min with constant mixing. Unbound cGAMP was recovered from the liquid phase, and the concentration was then assayed by mixing 4 µl of the solution with 2 µl CapV_Vc_-His_6_ (9.5 µM in FP-Rh buffer) as described above. Untreated cGAMP was used as an input control. Three technical repeats were included for each sample.

To determine if Gp15 interferes with cGAMP binding of CapV, an alternative cGAMP binding assay was used. Each cGAMP binding reaction (50 µl) contains 10 µl of purified CapV (9.5 µM) or FP-Rh buffer, 10 µl of Gp15 or BSA (each 10 µM in Gp15 dialysis buffer), 5 µl of 3’3’-cGAMP (5 µM), and 25 µl FP-Rh buffer. The reactions were incubated at RT for 30 minutes, and then filtered through Cytiva Vivaspin 500 10 Kda MWCO filters (Cytiva) at 12,000 g for 30 minutes. About 25 µl of filtrate was collected. Four µl of each filtrate was used in the FP-Rh assay by mixing with 2 µl CapV_Vc_-His_6_ (9.5 µM in FP-Rh buffer) as described above. Three technical repeats were included for each sample.

### LC-MS/MS quantification of cGAMP in solution

3ˈ3ˈcGAMP was measured using UPLC MS/MS as described previously [28]. The cGAMP samples harvested after the binding assay were first dried and then resuspended in 50 µL of ultra-pure water and 5 µl of this solution was injected into the UPLC-MS/MS system (Waters, Xevo). Electrospray ionization using multiple reaction monitoring in negative-ion mode at m/z 673.24→343.93 was used to detect 3ˈ3ˈcGAMP. The MS parameters were as follows: capillary voltage, 3.5 kV; cone voltage, 50 V; collision energy, 34 V; source temperature, 110 °C; desolvation temperature, 350 °C; cone gas flow (nitrogen), 50 L/h; desolvation gas flow (nitrogen), 800 L/h; collision gas flow (nitrogen), 0.15 mL/min; and multiplier voltage, 650 V. Waters BEH C18 2.1 × 50 mm column with a flow rate of 0.3 mL/min with the following gradient was used: solvent A (10 mM tributylamine plus 15 mM acetic acid in 97:3 water:methanol) to solvent B (methanol): t = 0 min; A-99%:B1%, t = 2.5 min; A-80%:B-20%, t = 7.0 min; A-35%:B-65%, t = 7.5 min; A-5%:B-95%, t = 9.01 min; A-99%:B-1%, t = 10 min (end of gradient). 125, 62.5, 31.25, 15.62, 7.81, 3.625, 1.9 nM of 3ˈ3ˈcGAMP (Biolog) was used to generate a standard curve.

### cGAMP concentration in cells

For intracellular measurements, overnight cultures of *V. cholerae* Δ*capV* Δ*dncV* carrying different plasmids were diluted 1000-fold in LB with Kan and Carb, grown for 2 hours, and then the cultures were induced with 100 µM IPTG along with 100 µg/ml of sulfamethoxazole and grown for another 4 hours. One ml of cells was pelleted by centrifugation (3 min, 14000 rpm), the supernatant was removed and resuspended in 100 µL ice-cold nucleotide extraction buffer [acetonitrile, methanol, ultra-pure water, formic acid (2:2:1:0.02, v/v/v/v)]. The tubes were incubated at −20°C for 30 minutes, pelleted by centrifugation (5 min, 14000 rpm), and supernatants were transferred to a new tube and dried for 2 hours in a speedvac concentrator. Dried nucleotides were dissolved in 100 µL ultra-pure water and 5 µl of this solution was injected into the UPLC-MS/MS system (Waters, Xevo) as described above. The intracellular concentrations of 3’3’cGAMP were determined using a modified method previously published [79].

### Bacterial two-hybrid assay

Bacterial adenylate cyclase two-hybrid (BACTH) assays were performed using *E. coli* BTH101 cells following the manufacturer’s protocol (Euromedex). The *gp15* gene of ICP3 was cloned into pUT18 and pKNT25 plasmids, while *capV* of *V. cholerae* was cloned into the pKNT25 plasmid. These plasmids were maintained in DH5α. Fifty nanograms of each T18 or T25 plasmid constructs were co-transformed into 50 µl of chemically competent BTH101 cells. Empty vectors of all plasmids mentioned were used as negative controls in the co-transformation experiments. A portion of each transformation was plated onto LB agar containing 50 µg/ml kanamycin, 50 µg/ml carbenicillin, 100 µM IPTG, and 40 µg/ml X-gal. Plates were incubated at 37°C for 17 hours, followed by 24 hours at 30°C. The color of the colonies was visually inspected. β-galactosidase assay was performed to quantify *lacZ* gene expression in these transformants.

### CBASS gene expression assay

*V. cholerae* strain carrying a P*_capV_*-*luxCDABE* luciferase reporter plasmid was used to measure gene expression of the CBASS operon [51]. Single colonies were inoculated into a black 96-well plate with a clear bottom with 150 µl LB containing 50 µg/ml Kan and 2.5 µg/ml Cm in each well. Mineral oil (40 µl) was added to each well and incubated in a plate reader with constant shaking at 30°C for 20 h. OD_600_ and luminescence were recorded every 30 minutes.

### Quantitative Western blot to determine CapV and Gp15 cellular concentration

Anti-Gp15 and Anti-CapV antibodies were raised in rabbits and rats, respectively, by Pocono Rabbit Farm using recombinant Gp15_ICP3_ and CapV_Vc_ proteins. *V. cholerae* (WN6145) was infected with ICP3 at an MOI of 5 as previously described [52], and 1 ml of infected cells was collected at different time points (0 to 25 mins post-infection). A portion of these samples was used for viable cell count by serial dilutions and plating. The remaining cells were harvested by centrifugation immediately after sampling and suspended in 20 µl SDS-sample buffer (67.6mM Tris-HCl, pH 6.8, 3% SDS, 10% Glycerol, 5% β-mercaptoethanol, 0.05% Bromophenol Blue). Samples were heated at 95 °C for 5 minutes, chilled on ice for 2 minutes, centrifuged for 5 minutes, and 10 µl of the supernatant was used for SDS-PAGE electrophoresis. For the Gp15 standard curve, 4.8, 2.4, 1.2, and 0.6 pmol of purified Gp15_ICP3_ was mixed with 10 µl *V. cholerae* lysate prepared from WN6145 as described above. After transferring the proteins onto a 0.45 µm nitrocellulose membrane, the membrane was blocked with Everyblot blocking buffer (Bio-Rad), then incubated with anti-Gp15 serum (1:500 dilution in 1× TBST with 3% BSA) for 1 hour. The membrane was then washed 3 times in TBST (5 minutes each), followed by incubation with Goat anti-rabbit IgG, DyLight 680 (ThermoFisher), diluted 1:10000 in TBST with 3% BSA for 1 hour. The membrane was washed 3 times in TBST, followed by imaging with Li-Cor Odyssey CLx imager. The fluorescence intensity was measured using ImageJ. The amount of Gp15 was calculated from the standard curve and the copy number per cell was normalized to the MW of Gp15 monomer and the viable cell count in each sample. To determine the CapV amount in the cell, similar procedures were used to prepare cell lysates from *V. cholerae* strains without phage infection. For the CapV standard curve, 4.8, 2.4, 1.2, 0.6, and 0.3 pmol of purified CapV_Vc_ was mixed with 10 µl *V. cholerae* lysate prepared from WN6031 (Δ*capV* Δ*dncV*). Strain WN5735 was used for the CapV overexpression sample. Primary Rat anti-CapV serum was diluted 1:1000 in 1X TBST with 3% BSA before use, and the secondary antibody was Goat anti-Rat IgG (H+L) Cross-Adsorbed Secondary Antibody, DyLight™ 800 (ThermoFisher) diluted 1:10000 fold in TBST with 3% BSA. Blocking, washing, detection, and CapV copy number calculation were performed as above.

## Acknowledgments

We thank Dr. Aaron Whiteley for providing some bacterial strains and plasmids for this study. We thank Dr. Aimee Shen for providing fluorophosphonate-rhodamine. We thank the Ng and Camilli lab members for their insightful comments and discussions.

## Supporting information Captions

**Supplementary Table S1**: The effect of overexpression of ICP3 phage genes in changing *V. cholerae* El Tor SMX sensitivity.

**Supplementary Fig. S1**. Expression of *gp15* does not reduce expression of the CBASS operon in *V. cholerae*. The P*_capV_*-*luxCDABE* was used to measure the gene expression of the CBASS operon (*capV-dncV-cap2-cap3*) in *V. cholerae* strains with and without ectopic expression of *gp15* of ICP3. Light production was normalized to OD_600_.

**Supplementary Fig. S2**. (A) Amino acid similarity and sequence alignment of Gp15 from ICP3, T7, and Mak (Alignment was performed with Blosum62 cost matrix with threshold 0). (B) T2 plaquing efficiency on different *E. coli* strains with and without CBASS_EC_ and GP15_T7_. The experiment was repeated more than three times, and representative results are shown.

**Supplementary Fig S3**. (A) Multiple repeats of Western blot analysis using anti-Gp15 antibodies with cell lysates prepared at different time points after ICP3 infection of *V. cholerae* WT. (B) Standard curves of Western blot signals (fluorescence) from different concentrations of purified Gp15 and CapV mixed with *V. cholerae* lysates lacking these proteins. Each experiment was repeated three times, and the standard curve generated from all trials are shown.

**Supplementary Table S2: Strains and plasmids used in this study.**

**Supplementary Table S3**: primers used in this study.

